# Two-photon imaging of excitatory and inhibitory neural response to infrared neural stimulation

**DOI:** 10.1101/2024.02.28.582632

**Authors:** Peng Fu, Yin Liu, Liang Zhu, Mengqi Wang, Yuan Yu, Fen Yang, Weijie Zhang, Hequn Zhang, Shy Shoham, Anna Wang Roe, Wang Xi

**Affiliations:** Interdisciplinary Institute of Neuroscience and Technology (ZIINT), the Second Affiliated Hospital, School of Medicine, Zhejiang University, Hangzhou, 310020, China; Key Laboratory of Biomedical Engineering of Ministry of Education, College of Biomedical Engineering and Instrument Science, Zhejiang University, Hangzhou, 310027, China; MOE Frontier Science Center for Brain Research and Brain Machine Integration, Zhejiang University, Hangzhou, 310058, China; Department of Ophthalmology and Tech4Health and Neuroscience Institutes, NYU Langone Health, New York, NY 10016, USA; Laboratory for Neuro- and Psychophysiology, Department of Neurosciences, KU Leuven Medical School, Leuven, 3000, Belgium

**Keywords:** infrared neural stimulation (INS), cell types, population response, neurostimulation

## Abstract

**Significance:** Pulsed infrared neural stimulation (INS, 1875 nm) is an emerging neurostimulation technology that delivers focal pulsed heat to activate functionally specific mesoscale networks and holds promise for clinical application. However, little is known about its effect on excitatory and inhibitory cell types in cerebral cortex.

**Aim:** Estimates of summed population neuronal response timecourses provide a potential basis for neural and hemodynamic signals described in other studies.

**Approach:** Using two-photon calcium imaging in mouse somatosensory cortex, we have examined the effect of INS pulse train application on hSyn neurons and mDlx neurons tagged with GCaMP6s.

**Results:** We find that, in anesthetized mice, each INS pulse train reliably induces robust response in hSyn neurons exhibiting positive going responses. Surprisingly, mDlx neurons exhibit negative going responses. Quantification using the index of correlation illustrates responses are reproducible, intensity-dependent, and distance-dependent. Also, a contralateral activation is observed when INS application.

**Conclusions:** In sum, the population of neurons stimulated by INS includes both hSyn and mDlx neurons; within a range of stimulation intensities, this leads to overall excitation in the stimulated population, leading to the previously observed activations at distant post-synaptic sites.

## Introduction

Understanding complex neural circuits and their relationship to specific behaviors involves precise temporal and spatial modulation of neuronal subtypes. Non-genetic near-infrared optical stimulation was one of the promising non-invasive neural interface technologies for the brain ^1-5^. Recently, pulsed infrared neural stimulation (INS) technique has been introduced as a method capable of modulating neural activity safely and reversibly ^1^. In contrast to effects induced by alternative wavelengths of infrared stimulation (e.g., 808 nm ^2^, 980 nm ^3^, 5.6 μm ^4,5^), the pulsed delivery of ∼1.875 μm infrared wavelengths leads to a focal delivery of heat and rapid absorption by water ^6^. When delivered via 200 μm fiber optic in short pulse trains (0.25 ms, 200 Hz, 0.5 s), this highly focal (submillimeter) optical approach has provided a unique way of functional column-specific stimulation in primate cortex ^7^. Thus, the advantages of INS over conventional electrical stimulation include high spatial selectivity, contactless delivery, and importantly for primate and human application, neuromodulation of brain sites without prior opsin expression ^8,9^. Furthermore, with this precision of targeted optical fiber stimulation and MRI compatibility, focal INS combined with MRI can be used for *in vivo* mapping of brain networks in primates ^10-12^ and holds promise for neuromodulation in awake behaving monkeys. While such applications have demonstrated great promise for circuit neuromodulation *in vivo*, its mechanism of action or effect on individual cell types is currently still poorly understood.

There is now a growing body of evidence showing that INS leads to neuromodulation. INS has been shown to induce excitatory and inhibitory neuronal responses in anesthetized rodents, as assessed with electrophysiology, intrinsic signal optical imaging, and calcium imaging *in vivo* ^13,14^. INS stimulation of visual cortex in anesthetized macaque monkeys produced responses typical of visually induced cortical intrinsic signals ^7^ and furthermore led to selective modulation of functionally matched ocular dominance domains, consistent with activation of local cortico-cortical connections. The demonstration that INS stimulation in ultrahigh-field MRI could lead to activation of anatomically predicted mesoscale global brain sites in the macaque monkey further suggested that projection cells (excitatory pyramidal neurons) are being activated by INS ^10-12^. These INS-induced responses have been shown to be intensity- and duration-dependent.

Despite this compelling evidence, it has been challenging to directly demonstrate neuronal response electrophysiologically. One issue, referred as the Becquerel effect, is that the direct heating of the recording electrode contaminates the neuronal response with heat-induced current in the electrode. Cayce et al. used calcium imaging *in vivo* with simultaneous INS in anesthetized rodents and observed intracellular calcium signals in cortical astrocytes and apical dendrites on the brain surface ^14^. Kaszas et al. used two-photon calcium imaging with the genetically encoded calcium indicator Syn-GCaMP6f and showed that INS induces weak intracellular calcium signals in neurons in anesthetized mouse cortex *in vivo* ^15^. To date, our understanding of neuronal response is still rudimentary. The underlying mechanism of action is unknown ^16-23^ and the effect on responses of different neuronal subtypes as well as different physiological states *in vivo* at a cellular level is still lacking. In particular, even though INS has been shown to induce BOLD activation at distant cortical sites in fMRI studies, little is known about the cellular circuits contributions to this functional connectivity result.

To address how INS affects single neurons *in vivo* and to examine effects on different cell subtypes, we conducted two-photon imaging of neuronal calcium response to INS at single-cell resolution in layers 2/3 of mouse somatosensory cortex. Calcium response of hSyn- and mDlx-labeled neuronal subtypes was examined using specific genetically encoded calcium indicators GCaMP6s. We found that INS induced robust, intensity-dependent modulation of neuronal calcium reflectance change which occurred precisely in phase with the frequency of pulse train repetitions. In anesthetized mouse, hSyn neurons exhibited positively deflecting responses to INS. Surprisingly, mDlx neuron population contained varied responses, some of which exhibited negative going response, and may reflect diversity in the inhibitory neuron population. Thus, these data establish the effectiveness of INS on both hSyn and mDlx neurons, and possible dependence on cell subtypes. The implications of this finding are discussed.

## Materials and Methods

### Animals

All experimental animals were approved by the Zhejiang University Animal Experimentation Committee and were completed in compliance with the National Institutes of Health Guide for the Care and Use of Laboratory Animals. Female C57BL/6J mice (8-10 weeks old) were group-housed (3 per cage) on a 12 h light-dark cycle and provided with food and water *ad libitum*.

### Virus injection

For hSyn neurons imaging, we injected 200 nL rAVV2/9-hSyn-GCaMP6s-WPRE-hGH-pA (titer = 1.03×10^13^ vg/ml, PT-0145, BrainVTA) per site; for mDlx neurons imaging, we injected 200 nL rAAV2/9-mDlx-GCaMP6s-WPRE-hGH-pA (titer = 5.48×10^12^ vg/ml, PT-2757, BrainVTA) per site. For co-localization experiment, we injected 200 nL rAAV2/9-mDlx-GCaMP6s-WPRE-hGH-pA (titer = 5.48×10^12^ vg/ml, PT-2757, BrainVTA) and 200 nL rAAV2/9-hSyn-NES-jRGECO1a-WPRE-hGH-pA (titer = 5.82×10^12^ vg/ml, PT-1593, BrainVTA), respectively.

### Cranial window

Mice were anesthetized with isoflurane (5% inhalation, mixed with fresh air, 0.5 L/min) and placed in a stereotactic frame, after which isoflurane was maintained at 2% throughout the surgical procedure. The body temperature was maintained at 37 °C by a heating pad during the procedure. A craniotomy (measuring 4 mm in diameter, around the coordinate of 0.3 mm posterior and 2.3 mm lateral from bregma) was performed exposing the right somatosensory cortex using a dental drill. The dura was carefully removed and artificial cerebrospinal fluid (ACSF) was used to keep the brain tissue clean and viable. Then, we injected 200 nL of virus per site around the craniotomy center at 400 μm below the pia. About 3-5 sites of viruses were manually injected by a patch pipette (tip diameter of approximately 15-20 μm). A cover glass (measuring 6 mm in diameter) was used to cover the brain and secured with medical glue at its edge. A custom-made head-plate with screw holes was glued to the skull and the gap between the head-plate and the skull was secured with dental acrylic. A chronic imaging window was completed which can be used repeatedly immobilized in the same position in the two-photon microscopy. Mice received an intraperitoneal injection of ceftriaxone (2.5 mg/kg) and a subcutaneous injection of buprenorphine (0.05 mg/kg) for up to three days post-surgery and were allowed to recover for at least three weeks before they were imaged.

### Two-photon imaging

Mice were imaged using customized two-photon microscopy (2P PLUS, Bruker Corporation) coupled with a femtosecond mode-locked Ti: Sapphire laser (80 MHz, 140 fs, Chameleon Ultra II, Coherent Inc.) with 920 nm (for GCaMP6s only) or 965 nm (for GCaMP6s and jRGECO1a). A Pockels Cell (EO-PC, Thorlabs Corporation) was used to regulate laser power. For neuron imaging, power after the 4X objective (N4X-PF, 0.13 NA, Nikon) and 16X objective (N16XLWD-PF, 0.8 NA, Nikon) was limited to a maximum of 40 mW, dependent on depth. Emission light was filtered using a bandpass 525/70 filter for GCaMP6s and a bandpass 595/50 filter for jRGECO1a, detected by two GaAsP PMTs (model H10770, Hamamatsu Photonics). Images were collected at ∼30 Hz using a resonant-galvo scanner with 512×512 pixel resolution for functional imaging. Images were collected using a galvo-galvo scanner with 1024×1024 pixel resolution for structure imaging. For two-photon imaging in anesthetized experiments, isoflurane (0.6%, mixed with fresh air, 0.5 L/min) was used to anesthetize mice during imaging. For two-photon imaging in awake experiments, three days of routine handling of the mice to acclimate them to the imaging system and immobilization device, which greatly helped to reduce animal motion.

### Infrared neural stimulation

For all INS experiments, a single wavelength laser (FC-W series, CNI) coupled with multimode fiber (MM200, Newdoon; 200 μm core diameter, 0.37 NA) was used to provide an 1875 nm infrared beam. The laser fiber was placed into a custom-built micromanipulator and under two-photon microscopy. Non-damaging laser pulses (wavelength, 1875 nm; peak radiant exposure, from 0.1 to 1 J/cm^2^ per pulse at fiber tip) were calibrated with a power meter (PM320E, Thorlabs Corporation; coupling with thermal power sensor, S401C, Thorlabs Corporation) before every experiment and transmitted through an optical fiber positioned at a ∼45° angle to the brain surface. For multiple pulse train stimulation, each pulse train contains 6 bursts of 0.5 s duration (0.25 ms pulse width per pulse at 200 Hz, 100 pulses) and 2.5 s interval presented once every 60 s. In a single trial, the first 10 s was the baseline, following 18 s INS application and 32 s convalescence time. The laser powers were presented in randomized order to prevent trends. For long-lasting pulse train stimulation, pulse train contains 2 s duration (0.25 ms pulse width per pulse at 200 Hz, 400 pulses) once every 30 s. In a single trial, the first 3 s was the baseline, following 2 s INS application and 25 s convalescence time. The two-photon system was used to initiate a TTL signal, sending it to the laser to synchronize the infrared neural modulation and imaging.

### Monte Carlo simulation for infrared light

To simulate light propagation in planar multi-layered tissues, we modeled tissues in a cube of 4 mm slide length. A 4×4×4 mm cube of tissues in Cartesian coordinates was generated, discretizing space in 10 μm steps, which we modified the codes for our infrared light (λ = 1875 nm) propagation ^24,25^. The infrared light propagation emitted from the optical fiber (200 μm core diameter, 0.37 NA) and recording of simulated quantities were performed through three-layered tissue structures (the first layer is ACSF; the second layer is glass; the third layer is cortex) with a Cartesian-grid voxel-based method, which allows assignment of optical properties, including absorption coefficient (μ_a_ [mm^−1^]), scattering coefficient (μ_s_ [mm^−1^]), and anisotropy of scattering (g [dimensionless]) to each individual voxel. The current simulation code was fully implemented with MATLAB (version R2020b, MathWorks).

### Data analysis

Acquired time-lapse two-photon images were read and analyzed in FIJI (Fiji Is Just ImageJ, NIH), Cellpose (version 2.0) and MATLAB (version R2020b, MathWorks). Movement artifacts in the xy plane were corrected by using the template_matching plugin in FIJI ^26,27^. Cellpose was used to detect neuron soma except for neuropil and perform cellular segmentation to get masks ^28^. FIJI registers raw movies, uses masks, and extracts calcium fluorescence traces. Fluorescence changes (F) were calculated by averaging the corresponding pixel values in each specified ROI mask. In the following analysis, we used our self-written MATLAB codes. Relative fluorescence changes ΔF/F_0_ = (F-F_0_)/F_0_ were calculated as calcium transient, where the baseline fluorescence F_0_ was estimated as the first several seconds before stimulation of the entire fluorescence recording. For typical example in Figs. 2(a), 3(d), 3(e), 5(a) and 6(a), Kalman filter ^29^ was used to smooth the data in MATLAB.

For multiple pulse train stimulation, two special indices were used to evaluate the averaged response caused by the INS for each neuron. First, the correlation value was defined as a positive sine function (Y = sin 1/3×2π×X), whose frequency is 1/3 Hz, correlated between stimuli patterns, and INS-evoked neuronal response curve. The positive sine function was used to fit the oscillation induced by the six-consecutive pulse trains. For each radiant exposure, during the 10-28 seconds INS period, we calculated the correlation between the sine function and neuronal calcium response curve. A positive calcium response (excitatory effect by INS) was defined by the correlation value > 0.2. A negative calcium response (inhibitory effect by INS) was defined by the correlation value < -0.2. Second, the averaged peak-to-peak amplitude of calcium transient (ΔF/F_0_) was used as an index of neuronal response during the INS stimuli for distance assessment. We calculated six peaks during six trains and averaged these six amplitude values as amplitude. When correlation value was negative, its amplitude was negative and means a decrease induced by INS. When correlation value was positive, its amplitude was positive and means an increase induced by INS. For long-lasting pulse train stimulation, the total peak area under the curve during the 3-10 seconds period was measured as a stimulated effect. This value is affected by several choices in the analysis dialog in Prism: the definition of baseline (baseline = 0) and the definition of peaks too small to count (ignore peaks that are less than 10% of the distance from minimum and maximum).

### Statistical analysis

All data are presented as means ± SEM. A paired Wilcoxon test was performed to estimate the statistical differences in Fig. 4(f) and Fig. S3. An unpaired Mann-Whitney test was performed to estimate the statistical differences in Fig. S2. The comparison was analyzed using the Wilcoxon test when they can be paired one by one; otherwise, a Mann-Whitney test was used as indicated. In all tests, p < 0.05 is considered statistically significant and unmarked represents no significance. All statistical analyses were performed in Prism (version 8.0, GraphPad Software). No additional methods were used to determine whether the data met the assumptions of the statistical approach. The statistical details of each experiment can be found in the figure legends and tables.

## Results

We measured the INS-evoked *in vivo* calcium activity of populations comprising hSyn vs. mDlx neuronal subtypes expressing the hSyn-GCaMP6s and mDlx-GCaMP6s, respectively ^30^, as well as a two-color multi-labeling approach. We applied a robust version of a common pulsed INS stimulation paradigm (pulse trains consisting of short pulses 0.25 ms, high frequency 200 Hz, short pulse train duration 0.2-0.5 s). Stimulation was applied in sets of 6 pulse trains to assess reliability of response (Fig. S1 showed correction of the applied laser energy levels for the glass).

### Viewing calcium activity of 2 types of neurons in mouse cortex *in vivo*

For our initial attempts to investigate the properties of pan-neuronal calcium activity to INS in mouse somatosensory cortex, we employed the viral labeling strategies that AAV-expressed GCaMP6s ^30,31^ under the pan-neuronal promoter human synapsin (hSyn) packaged in serotype 2/9 (AAV2/9-hSyn-GCaMP6s), which mainly produces robust expression of GCaMP in excitatory neurons ^30,32^ [Fig. 1(a)]. To further observe the specific inhibitory neurons responding to INS, we adopted AAV-expressed GCaMP6s under an enhancer (mDlx) which induces expression in GABAergic neurons across a wide range of species, including mice ^30,32,33^ [Fig. 1(b)]. To examine whether these populations are distinct, we co-labeled the two cell subtypes using hSyn-jRGECO1a and mDlx-GCaMP6s in the same field of view (FOV) in same mouse cortex [Fig. 1(c); Fig. S2 showed spontaneous activity variations in two subtypes]. This revealed very few co-labeled neurons [Fig. 1(d), 6.8 ± 1.8%. Total 383 neurons in n = 3 FOVs from 2 animals; 219 hSyn-labeled neurons, 189 mDlx-labeled neurons, 25 merged neurons], indicating the cell populations labeled in Fig. 1(a) and Fig. 1(b) were distinct populations.

**Fig.1.**
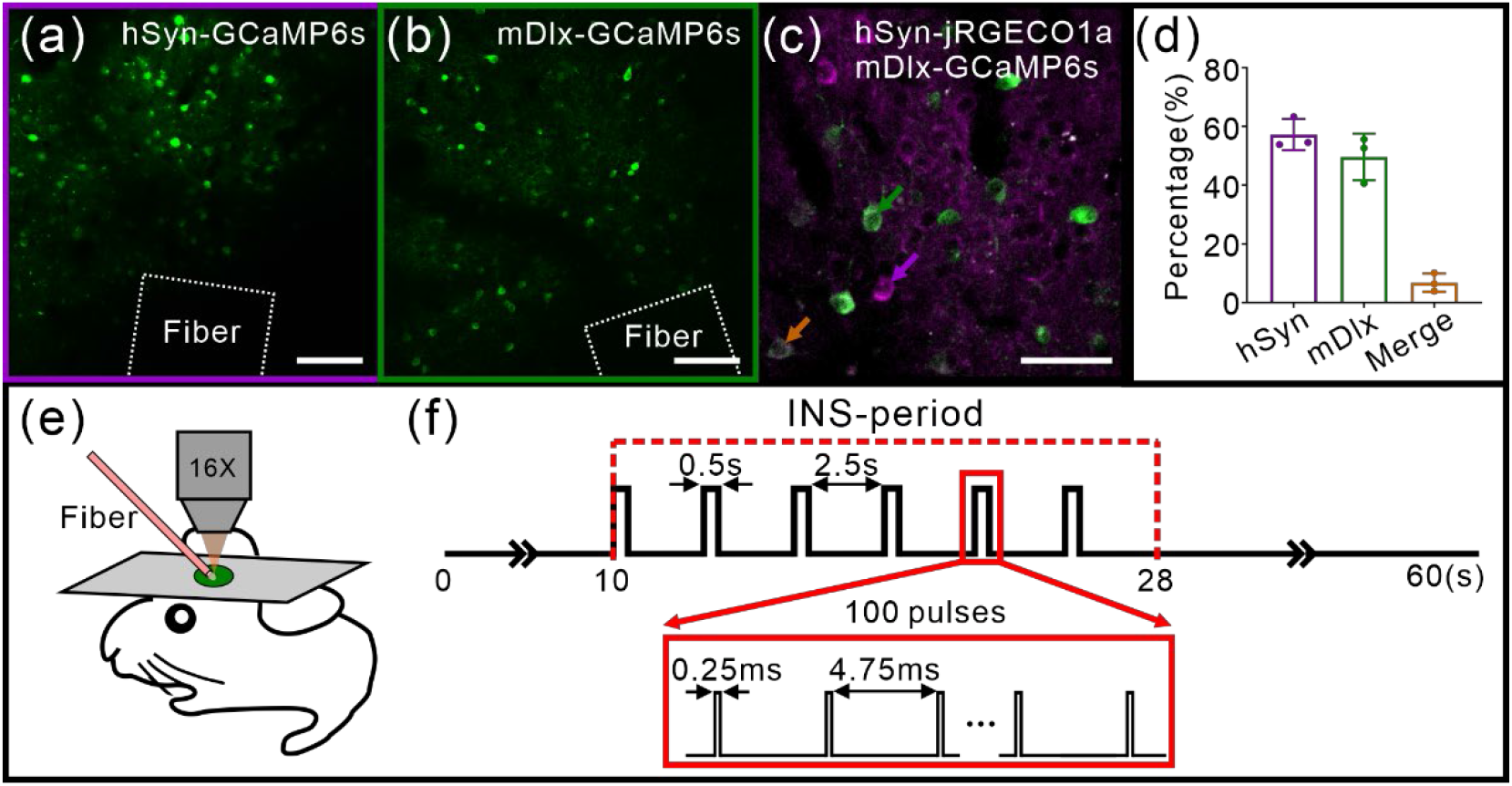
INS-TPM setup and specific subtypes calcium imaging of INS-induced neuronal response in mouse cortex. (**a**) hSyn-GCaMP6s expression in mouse somatosensory cortex labels hSyn neurons. The white dotted box indicates INS fiber tip position. Scale bar, 100 μm. (**b**) mDlx-GCaMP6s expression in mouse somatosensory cortex labels mDlx neurons. The white dotted box indicates INS fiber tip position. Scale bar, 100 μm. (**c**) hSyn-jRGECO1a (purple) and mDlx-GCaMP6s (green) co-expression in the same mouse somatosensory cortex labels different population neurons, respectively. Scale bar, 50 μm. (**d**) Proportion of hSyn-jRGECO1a (purple) and mDlx-GCaMP6s (green) expression neurons, respectively. Error bars are mean ± SEM. (**e**) INS-TPM setup, the mouse brain was fixed under the TPM and the somatosensory cortex was covered by a glass. The optical fiber (pink bar) touched on the cover glass for stimulation. (**f**) INS stimulation paradigm. One INS stimulation comprises six pulse trains and each train comprises 100 pulses at single energy. Train duration: 0.5 s, train interval: 2.5 s, repeat 6 times; pulse width: 0.25 ms, repetition rate: 200 Hz; radiant exposure: 0.16-0.79 J/cm^2^.

We employed two-photon calcium imaging to visualize these two subtypes of neuronal calcium activity during INS application separately. In these experiments, we applied a variant of previously established stimulation parameters that were shown to induce neuromodulation in a non-damaging fashion ^13,14,34^. In our previous studies ^7^, using a 200-μm optical fiber aimed at targeting 200-300 μm sized cortical columns, we used trains of very brief pulses (short pulse 0.25 ms, high frequency 200 Hz, short pulse train duration 0.5 s, radiant exposure 0.1-0.8 J/cm^2^ per pulse, ISI of 2.5 s); this delivered small boli of heat absorption (thermal confinement regime) to the tissue. Here, to better observe the reliability of response to INS in mouse [Fig. 1(e)], we modified the previous paradigm of 3-4 pulse trains ^11^ to 6 such pulse trains and recorded for 60 seconds per trial [Fig. 1(f)]. Radiant energies were non-damaging ^9,34^ and neurons maintained normal activity levels after INS application (cf., Fig. S3). Below, we present results from hSyn neurons (Figs. 2-4) and then from mDlx neurons (Fig. 5).

**Fig.2.**
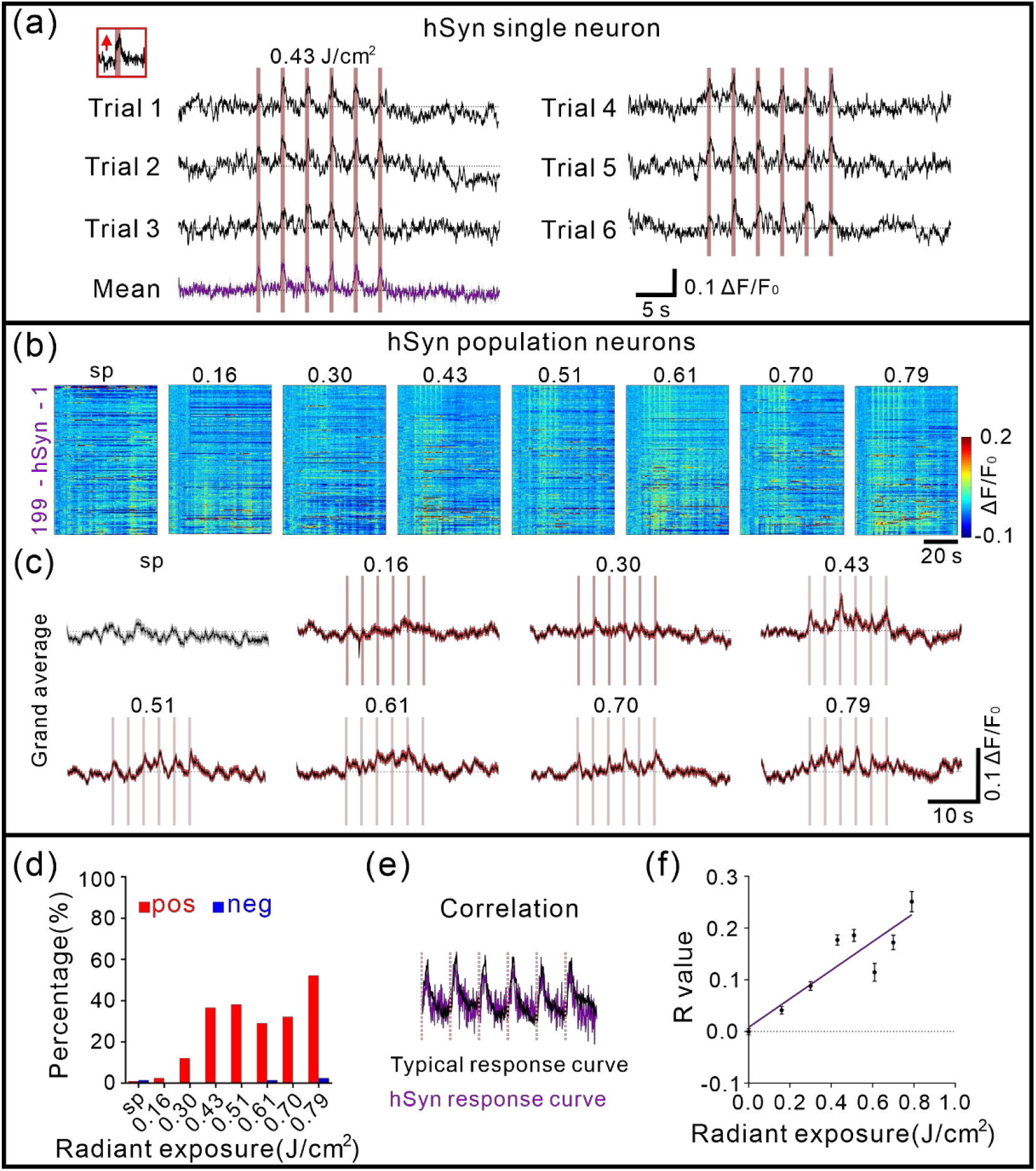
INS-induced calcium activity of hSyn neurons. (**a**) Positive response of a typical hSyn neuron to INS (radiant exposure: 0.43 J/cm^2^). The pink bar indicates the INS period. The insert panel indicated the enlarged time response during the first INS pulse train. Bottom purple curve was mean. (**b**) All trial-averaged responses of hSyn neurons (n = 199 neurons in 3 anesthetized mice) and a pseudo-colored plot summarizing the trial-integrated calcium activity trace for each neuron during INS intensity at 0.16, 0.30, 0.43, 0.51, 0.61, 0.70 and 0.79 J/cm^2^, sorted by correlation during the INS period, respectively. (**c**) A grand average of spontaneous and INS-induced calcium activity traces for all neurons corresponding to b. (**d**) Percentage of positive calcium activity induced by INS in hSyn neurons (red, 1.0%, 2.5%, 12.1%, 36.7%, 38.2%, 29.1%, 32.2% and 52.3% at 0, 0.16, 0.30, 0.43, 0.51, 0.61, 0.70 and 0.79 J/cm^2^, respectively). pos: positive response; neg: negative response. (**e**) Calculation of correlation index for hSyn neurons. (**f**) Correlation of calcium responses curve in hSyn Near field neurons (fitting curve, Y = 0.2755×X+0.0081, R^2^ = 0.8062, p = 0.0025). Data represents mean ± SEM in Table S1.

### INS induced calcium activity: hSyn neurons

Here, we exhibited the effects of INS on hSyn neurons in anesthetized mice. As shown in Fig. 2(a) with a single hSyn neuron, using real-time (∼30 Hz acquisition rate) two-photon imaging over the entire course of INS [1 trial total 60 s: 10 s baseline, 18 s INS period, 32 s recovery period, see Fig. 1(f)], we found that indeed the pulse trains (each indicated by pink vertical line) modulated neuronal calcium response (black traces) in phase with the pulse train delivery. Similar to a previous study ^15^, we observed positive deflections from baseline in the calcium signal responses of hSyn neurons [Fig. 2(a), red arrow in top left inset]. As shown in the 6 trials, which is clearly seen in the mean of 6 such trials [bottom purple trace in Fig. 2(a)], this was a consistent response across the six successive pulse trains in each trial and consistent across trials.

*In vivo* calcium responses were recorded in 199 hSyn neurons in anesthetized mice [n = 3 mice, Fig. 2(b)]. Previous studies characterizing INS have indicated that radiant exposure is a primary parameter for determining the strength of response ^7,10,13,15^. We delivered different laser radiant exposures of INS; these values (0.1-0.8 J/cm^2^) were guided by irradiance values determined to be non-damaging and effective in our previous studies ^10,11,13,14^. INS intensities were applied at radiant exposures of 0.16, 0.30, 0.43, 0.51, 0.61, 0.70 and 0.79 J/cm^2^ [INS intensity correction shown in Fig. S1]. Populations of hSyn neurons responded robustly to INS in anesthetized state [mean traces shown in Fig. 2(c)] and most of them exhibited the positive going behavior [n = 5, 24, 73, 76, 58, 64 and 104 out of 199 neurons at 7 intensities, respectively; percentage shown in Fig. 2(d)]. Thus, under anesthesia, hSyn neurons predominantly respond with reliable and reproducible positive deflections.

To understand the different aspects of neuronal behavior to INS, we quantified characteristic correlation between the standard response (averaged positive responses to INS at 0.79 J/cm^2^) and each response timecourse [see calculation paradigm in Fig. 2(e); for more details in Materials and Methods]. For hSyn Near field neurons (n = 140 neurons from 3 anesthetized mice. The definition is described below in Fig. 3; for Far field neurons, see Fig. S5), as shown in Fig. 2(f), correlation values exhibited, at the lower radiant exposure, a moderate correlation, while strong correlations emerged at higher energies. The fitted correlation curve showed increased correlation value with the increase in laser radiant exposures (linear fitting), illustrating clear intensity dependence in Fig. 2(f). Thus, by correlation measures the effect of increasing INS intensities, for hSyn neurons, led to increased calcium positive response. This characterizes the intensity-dependence and supports the reliability and reproducibility of INS-induced response in hSyn neurons.

**Fig.3.**
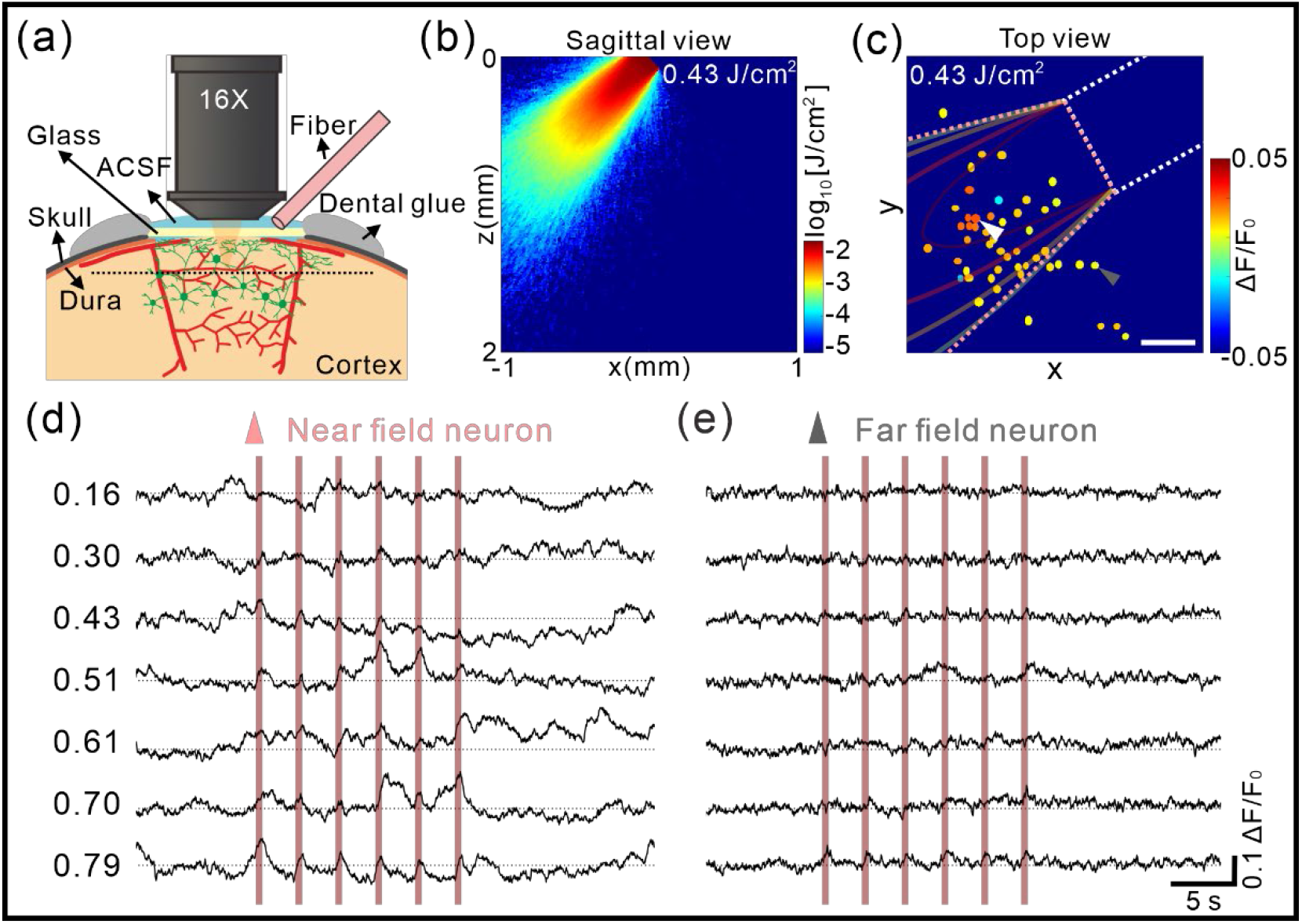
Focal effect of calcium responses to INS. (**a**) Illustration of the optical fiber placement and the imaging of calcium transients. An optical fiber tilted at an angle of ∼45° to the horizontal plane delivers infrared light to the cortex through the cover glass. (**b**) Theoretical infrared light distribution from a 200 μm core diameter fiber based on a Monte Carlo simulation. (**c**) Color coding of hSyn neurons spatial distribution map based on amplitude induced by INS. The white dotted box indicates fiber position. The pink dotted conoid indicates the light region. The red box indicates the field of view in two-photon imaging (Near field neuron: n = 40, pink triangle indicates the neuron in light direct exposure areas; Far field neuron: n = 9, gray triangle indicates the neuron outside light regions). Scale bar, 100 μm. (**d**-**e**), A representative example of INS-induced calcium transient in a Near (d, white triangle in c) and Far (e, gray triangle in c) field neuron over different laser radiant exposures, respectively. The pink bar indicates the INS period. Data (n = 6 trials) represents the mean.

### Focal effect of INS-induced calcium activity

To examine the spatial extent of INS, we conducted Monte Carlo simulation of the INS light distribution and examined neuronal responses inside and outside this region of INS illumination. These analyses were conducted based on our experimental setup [cartoon diagram shown in Fig. 3(a)]. Monte Carlo simulation was performed based on illumination with a 45° angle fiber to estimate the theoretical infrared light distribution [Fig. 3(b), for detail see Fig. S4]. In Fig. 3(c), the fiber position (shown as white dotted box) is shown in relation to all neurons in the field of view of two-photon imaging. Based on distance of neuron from the fiber tip center measurements (three-dimensional space distance), the relationship between amplitude and distance is hard to quantify. So, we next defined the light region based on Monte Carlo simulated light distribution (pink dotted trapezoid) and binned these neuronal responses into ‘Near’ vs. ‘Far’. After color coding each neuron by amplitude [see color scale in Fig. 3(c)], we found a distance-related effect across the population. Neurons within the light region are categorized as ‘Near’ field neurons [an example of hSyn neuron in Fig. 3(d), position shown with white triangle in Fig. 3(c)] and out of the light region as ‘Far’ field neurons [an example of hSyn neuron in Fig. 3(e), position shown with gray triangle in Fig. 3(c)]. The Near field neuron exhibits strong upward deflections (positive responses). The Far field neuron, in contrast, has a relatively weak response across all intensities tested. This suggests possible light intensity-related distance dependence. We further examined the calcium response to INS in contralateral cortex.

### INS induced calcium activity in the contralateral cortex

We anticipated seeing distally connected cortex’s activation response in addition to the local activation induced by INS, as shown in the previous fMRI study^10^. To address this distant response of neurons, we positioned the optical fiber as before and then observed the calcium activity in contralateral cortex [see setup of Figs. 4(a) and 4(b)]. Furthermore, we applied a long-lasting version of a common pulsed INS stimulation paradigm [400 pulses, as shown in Fig. 4(c)]. In the spontaneous state and the stimulation phase, we examined 335 hSyn neurons from 3 awake mice, respectively [as shown in Fig. 4(d), INS radiant exposure: 0.47 J/cm^2^]. A delayed peak is observed and lasted 5-10 seconds in the INS condition in Fig. 4(e). The activation response is shown by a quantification of the area under the curve following INS stimulation (during the 3-10 s period), and this allowed us to determine that the stimulation had an effect on the contralateral brain [see Fig. 4(f)]. This cellular population response can be considered to support the connection activation in the contralateral cortex of the previous INS-fMRI studies. Below we further investigate the calcium activity to INS in inhibitory neurons.

**Fig.4.**
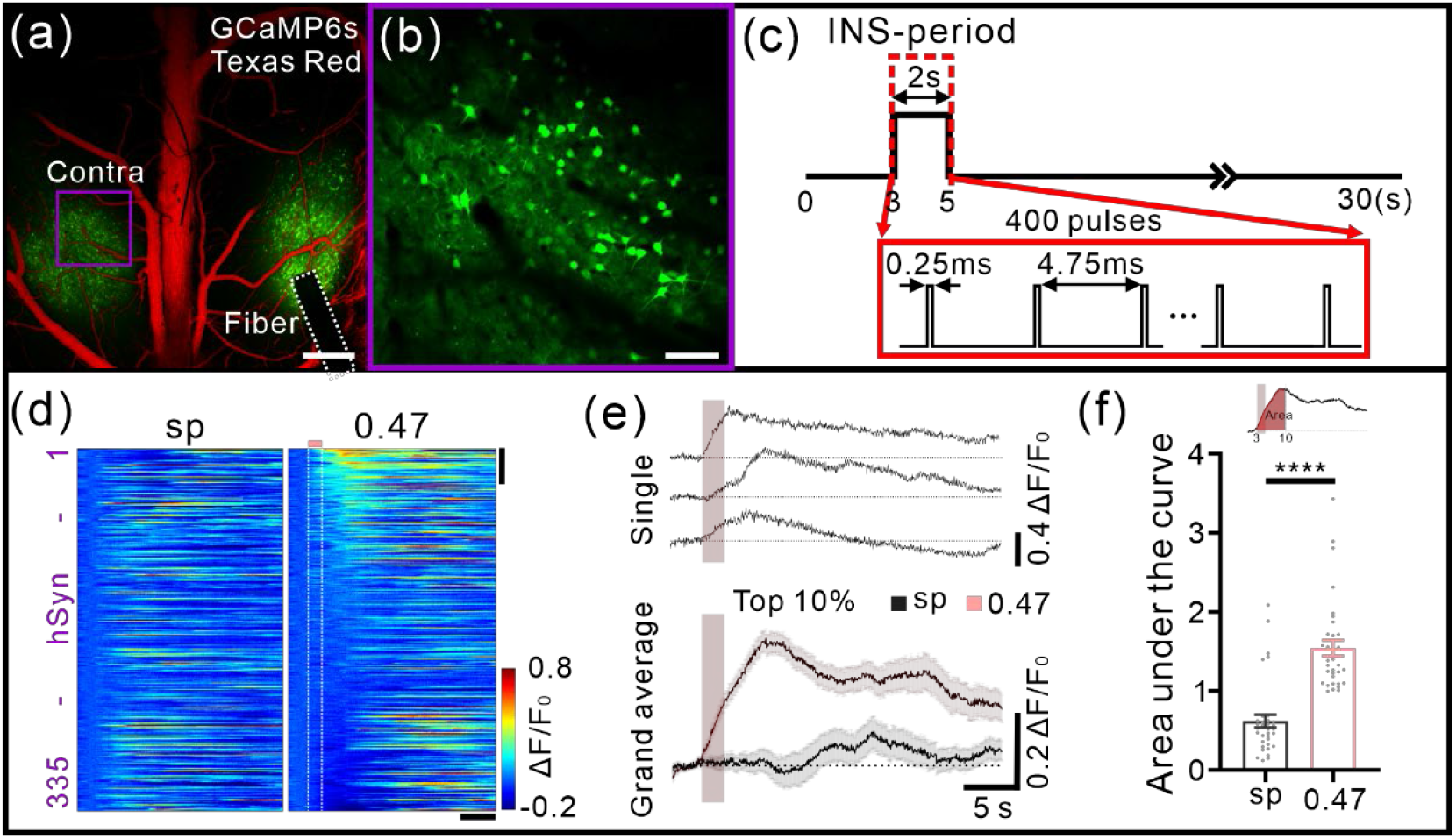
INS-induced calcium activity in the contralateral cortex. (**a**) Setup of contralateral imaging during INS stimulation. The white dotted box indicates INS fiber. The purple box indicates contralateral FOV. Scale bar, 500 μm. (**b**) hSyn-GCaMP6s expression in mouse contralateral cortex in a. Scale bar, 50 μm. (**c**) INS stimulation paradigm. One INS stimulation comprises a long-lasting pulse train (400 pulses). Train duration: 2 s; pulse width: 0.25 ms, repetition rate: 200 Hz; radiant exposure: 0.47 J/cm^2^. (**d**) All trial-averaged responses of hSyn neurons in the contralateral cortex (n = 335 neurons in 3 awake mice) and a pseudo-colored plot summarizing the trial-integrated calcium activity trace for each neuron during spontaneous baseline and INS, sorted by area under the curve during the 3-10 second period. (**e**) An example of INS-induced calcium activity traces and a grand average of top 10% neurons (n = 34) corresponding to d. (**f**) Area of calcium responses curve (during the 3-10 seconds period) in top 10% neurons between spontaneous baseline and INS-induced activity (area, sp: 0.6187 ± 0.0816, 0.47 J/cm^2^: 1.5440 ± 0.0988, p < 0.0001). Wilcoxon test; data represents mean ± SEM.

### INS induced calcium activity: mDlx neurons

In addition to study of hSyn neurons, we also examined responses of inhibitory (GABAergic) neurons labeled by mDlx promoter [an example of single mDlx neuron calcium response to INS shown in Fig. 5(a)]. Interestingly, in contrast to hSyn neurons, we find that many mDlx neurons, in anesthetized state, responded to INS with negative calcium fluorescence changes [see Figs. 5(b) to 5(d), n = 143 mDlx neurons from 3 mice]. Responses also appeared quite heterogeneous [evidenced by multiple examples of mDlx neuron responding to INS in Fig. 5(b) and the larger variability in response across neurons in Fig. 5(c), compared with Fig. 2(b)], perhaps due to diversity of GABAergic neurons ^35^. Out of 143 mDlx neurons recorded in anesthetized mice, using INS intensities at radiant exposures of 0.16, 0.30, 0.43, 0.51, 0.61, 0.70 and 0.79 J/cm^2^, up to 40% exhibited negatively going responses [Fig. 5(e)].

**Fig.5.**
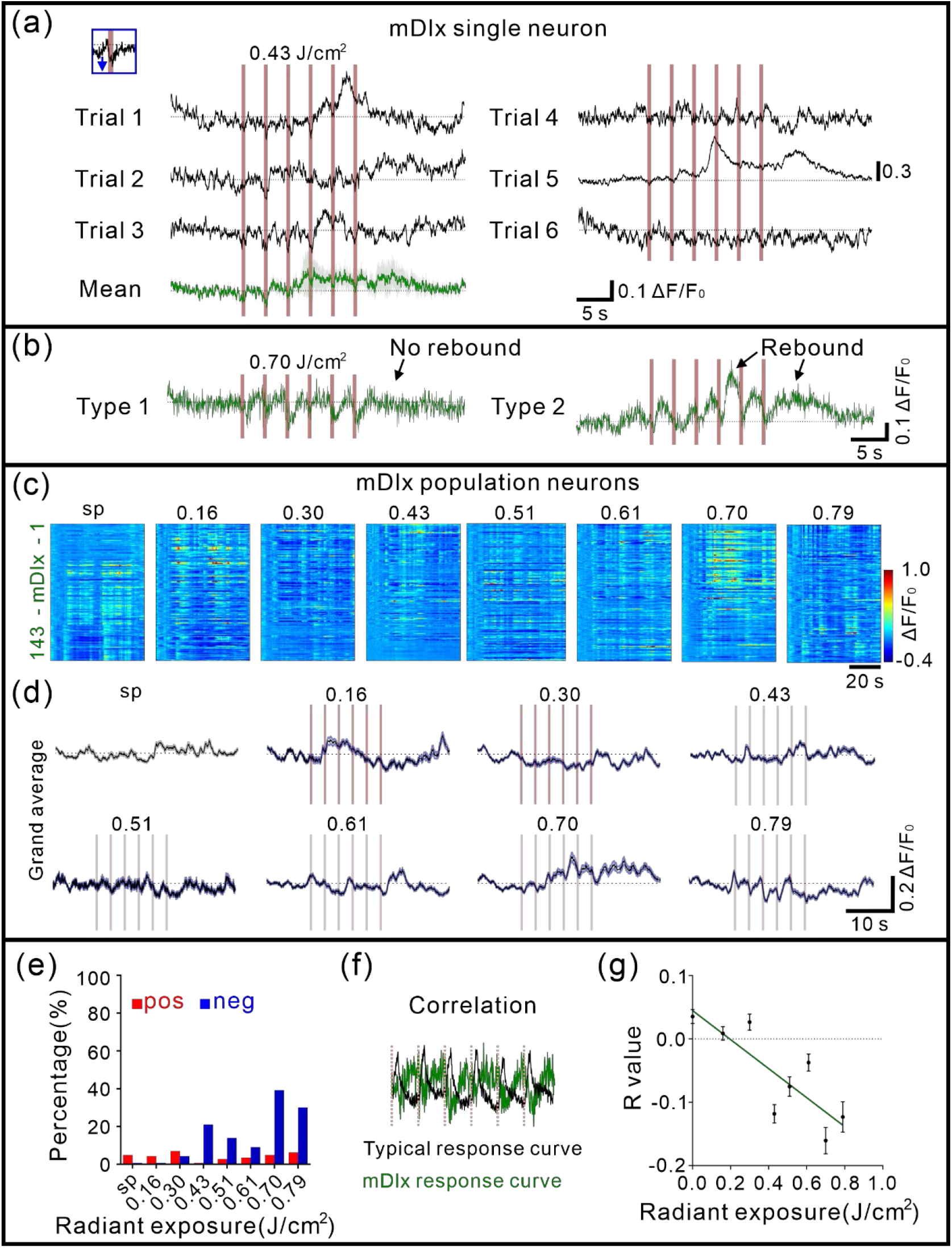
INS-induced calcium activity of mDlx neurons. (**a**) Negative response of a typical mDlx neuron to INS (radiant exposure: 0.43 J/cm^2^). The pink bar indicates the INS period. The insert panel indicated the enlarged time response during the first INS pulse train. Bottom green curves were mean. (**b**) Two types of negative responses of mDlx neurons. (**c**) All trial-averaged responses of mDlx neurons (n = 143 neurons in 3 anesthetized mice) and a pseudo-colored plot summarizing the trial-integrated calcium activity trace for each neuron during INS intensity at 0.16, 0.30, 0.43, 0.51, 0.61, 0.70 and 0.79 J/cm^2^, sorted by correlation during the INS period, respectively. (**d**) A grand average of spontaneous (gray) and INS-induced calcium activity traces for all neurons corresponding to b. (**e**) Percentage of negative calcium activity induced by INS in mDlx neurons (blue, 0.7%, 0.7%, 4.2%, 21.0%, 14.0%, 9.1%, 39.2% and 30.1% at 0, 0.16, 0.30, 0.43, 0.51, 0.61, 0.70 and 0.79 J/cm^2^). pos: positive response; neg: negative response. (**f**) Calculation of correlation index for mDlx neurons. (**g**) Correlation of calcium responses curve in mDlx Near field neurons (fitting curve, Y = - 0.2291×X+0.0447, R^2^ = 0.6869, p = 0.0110). Data represents mean ± SEM in Table S1.

For mDlx Near field neurons (n = 88 neurons from 3 anesthetized mice, for Far field neurons, see Fig. S5), we quantified correlation as defined above for INS-induced calcium activity [see calculation paradigm in Fig. 5(f)]. Index is negative, reflecting the negative reflectance changes in mDlx neurons. The fitted curves for correlation exhibited increasing negative values with increasing radiant exposures [Fig. 5(g)]. This intensity dependence was observed for correlation. Thus, for mDlx neurons, increasing INS intensities led to increased negative calcium responses during the pulse train delivery period. This signifies that the unusual negative calcium is not only reproducible but responsive to intensity in a predictable manner. At a population level, it appears that INS stimulation produces negative calcium transients in mDlx neurons.

These results suggest that, in both hSyn and mDlx neurons, neuronal calcium response to focal INS is observed primarily within the region of INS illumination ^14,15^. This further supports that neuronal response due to INS stimulation is quite focal and intensity-dependent.

## Discussion

### Summary

Previous studies have shown that pulsed infrared neural stimulation leads to neural response in the peripheral and central nervous system. However, the cellular basis of this response is not well understood. In this study, we examined the effect of INS stimulation on excitatory and inhibitory cortical neurons (hSyn-GCaMP6s and mDlx-GCaMP6s) using two-photon calcium imaging in mouse somatosensory cortex. Our results describe robust and reproducible calcium signals in single cells. However, responses to INS in excitatory and inhibitory neurons were very different in anesthetized mice. For hSyn neurons, our results describe robust and reproducible positive deflecting calcium signals in single cells. In contrast, for mDlx neurons, INS reliably induces a negative deflection during the light delivery. In sum, although the mechanism underlying INS effects on cellular response requires further study, our study definitively establishes that single neurons in cerebral cortex, both excitatory and inhibitory cortical neurons, are robustly and reproducibly induced by INS. This puts INS on more solid ground for applications in neuroscience, engineering, and medicine.

### Comparison with other INS studies

Previous studies have seen both excitatory and inhibitory effects of 1875 nm INS. Electrical single-unit and multi-unit recordings in peripheral nervous system ^36^, cochlea ^37^, and macaque and rat cerebral cortex ^7,13^ have revealed the presence of excitatory effects on neurons. Patch-clamp recordings were used to explicitly characterize infrared-evoked depolarising potentials and currents ^20,38^. Calcium imaging has also reported neuronal activation by INS, but single neuron calcium response remained little studied ^14,15^. Using optical intrinsic signal imaging and fMRI, robust and intensity-dependent INS-induced hemodynamic signals have been recorded not only at INS stimulation sites but also at distant connected sites ^7,10-13^, a finding suggesting robust population level pyramidal cell activation. There are also studies that point to inhibition evoked by pulsed INS. *In vitro*, studies revealed INS increased the amplitude and frequency of spontaneous inhibitory postsynaptic currents, and that this response is mediated by GABAergic neurotransmission ^39^. Infrared inhibition has also demonstrated block of peripheral nerve conduction, suppression of axonal excitability and downstream synaptic transmission ^23,40-44^. Here, we show that INS pulse trains can induce positive deflections and negative deflections in hSyn and mDlx neurons, respectively.

### Contribution to population response

We now consider what effect these observations have at the population level in anesthetized state. As INS is non-cell type specific, both excitatory and inhibitory neurons within the INS optical light field are impacted. To estimate what a population response might look like, we calculated the weighted sum of our two hSyn (purple trace) and mDlx (green trace) population traces, based on the standard ratio of 80% excitatory and 20% inhibitory neurons in cerebral cortex ^45^. As shown in Fig. 6(a) (see black trace), this population timecourse appears quite similar to the hSyn neuron responses. Prior to INS stimulation, the population maintains a baseline spontaneous activity [proportions of cells in each phase indicated in Fig. 6(b)] and then when infrared light is irradiated on excitatory (hSyn-labeled) and inhibitory (mDlx-labeled) neurons, the population is observed to oscillate in phase with the occurrence of pulse trains. This is then followed by a recovery response [Fig. 6(b), population activity model]. This population response, we suggest, is consistent with the presence of increased spiking activity during and following INS pulse train stimulation.

**Fig.6.**
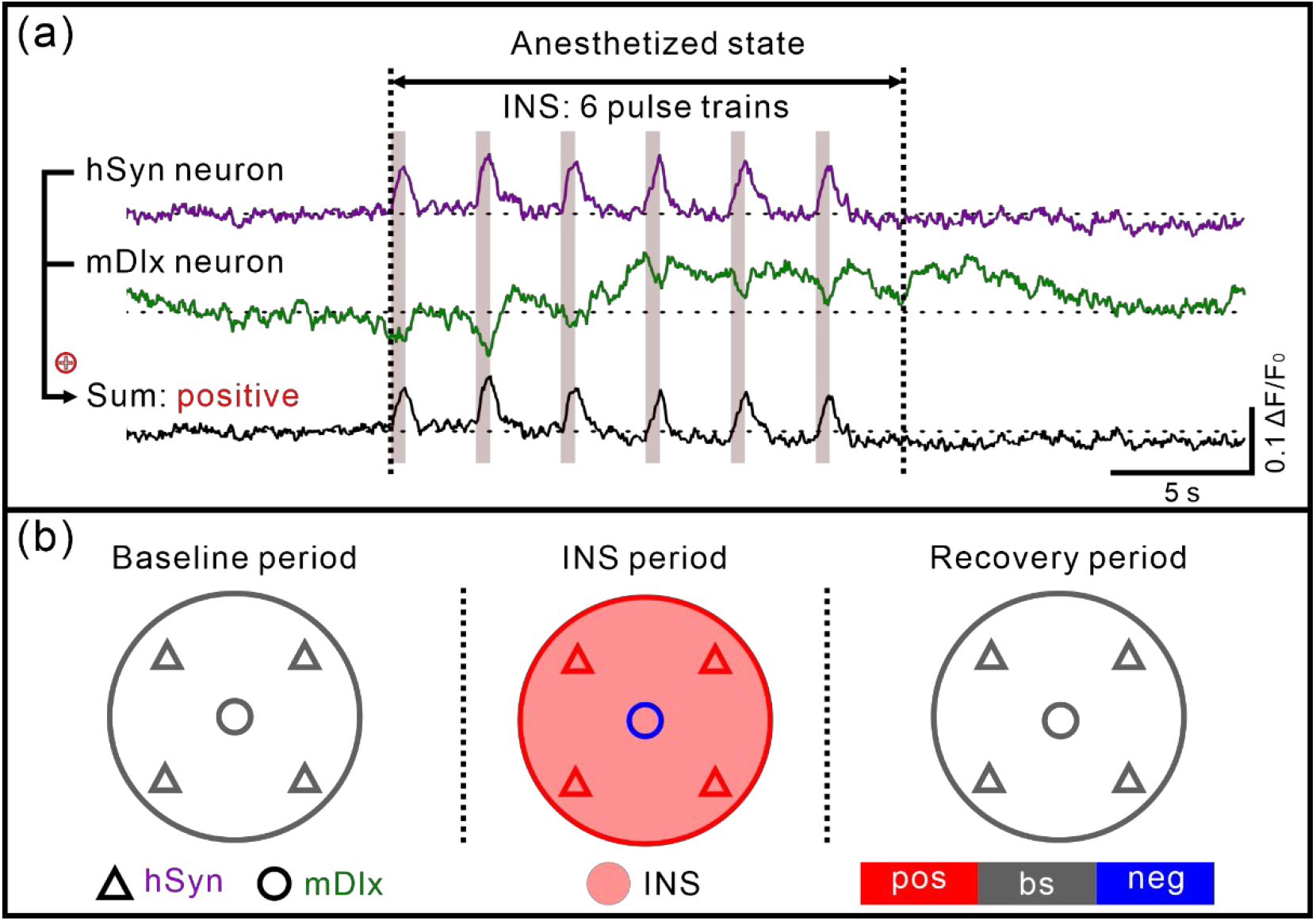
Modeling of INS-induced population neural calcium activity. (**a**) Timecourse of hSyn and mDlx neuron calcium response to INS (for example 0.43 J/cm^2^). The sum response consists of 80% hSyn response and 20% mDlx response (purple: hSyn, green: mDlx, Sum: black). The pink vertical bar indicates the INS period. (**b**). Population calcium response of regional hSyn neurons (triangle) and mDlx neurons (round) during different INS phases. The pink area indicates the INS irradiation region. pos: positive response (red); bs: spontaneous baseline (gray); neg: negative response (blue).

We further suggest that this population response is the basis for the hemodynamic responses consistently observed in INS-induced intrinsic optical imaging and BOLD fMRI studies. While the BOLD signal is a complex hemodynamic response which includes spiking and subthreshold cellular responses, as well as neuronally related and unrelated vascular contributions, the association of BOLD response with neuronal activity is generally accepted and has formed the basis for many fMRI studies. Based on a large number of intrinsic signal optical imaging studies and fMRI studies, neuronal activation leads to a brief ‘initial dip’ (seen optically as a small and reliable 0.1-1% decrease in reflectance, related to the deoxygenation hemodynamic phase, often not seen in fMRI due to small size, cf. reference ^46^), and is followed by a large positive BOLD signal (related to the influx of newly oxygenated blood) with a timecourse that lasts several seconds before returning to baseline. INS-induced BOLD signals have been observed both at the site of stimulation and at distant ‘functionally connected’ brain sites; these brainwide networks have been shown to be consistent with known anatomical networks ^11^ and have formed the basis for a mesoscale connectome project in macaque monkey ^10^. Note that the intensities used in these studies are typically lower (3-4 pulse trains, 0.1-0.3 J/cm^2^), directly delivered to the brain, and lead to quite focal columnar-scale stimulation. Here, we positioned the fiber with a 45° angle via a cover glass through a certain thickness of the cortex (depth we used in two-photon imaging). The actual intensity delivered to the neurons we observed is probably much lower. Thus, we suggest that there is sufficient excitatory neuron-driven activation (from the positive response period), to produce the hemodynamic signals seen robustly in BOLD fMRI, forming the basis for pyramidal cell mediated activation of functionally connected sites in fMRI studies.

## Code and Data Availability

Raw images and data are not publicly available at this time but may be obtained from the authors upon reasonable request.

## Credit Author Statement

Conceptualization, A.W.R., W.X., P.F., Y.L.; Methodology, P.F., Y.L., F.Y., W.X., A.W.R; Formal Analysis, P.F., Y.L., W.Z., W.X., A.W.R.; Investigation, P.F., Y.L., W.X., A.W.R., S.S.; Writing – Original Draft, P.F., W.X., A.W.R.; Writing – Review & Editing, P.F., W.X., A.W.R., Y.L., L.Z., M.W., Y.Y., W.Z., H.Z., S.S.; Visualization, P.F., W.X., A.W.R., Y.L., L.Z., M.W., W.Z., H.Z.; Supervision, W.X., A.W.R.; Project Administration, W.X., A.W.R., P.F., Y.L.; Funding Acquisition, W.X., A.W.R.

## Acknowledgments

We thank Y. Shu, L. Wang, X. Liu, and L.L. Wang for discussions and comments on the manuscript. This work was supported by the grants from National Key R&D program of China Brain Initiative 2021ZD0200401 (A.W.R; W.X.), 2018YFA0701400 (A.W.R); National Natural Science Foundation of China U20A20221 (A.W.R), 81961128029 (A.W.R), 31627802 (A.W.R); Key Research and Development Program of Zhejiang Province 2022C03096 (W.X.), 2020C03004 (A.W.R); Fundamental Research Funds for the Central Universities 226-2022-00083 (W.X.), 2019XZZX003-20 (A.W.R).

