## supplemental file for "Two-photon imaging of excitatory and inhibitory neural response to infrared neural stimulation"

### Supplemental Material

Supplemental Table 1 and Supplemental Figures 1-5.

**Table S1. Correlation values for Near and Far field in hSyn and mDlx neurons.**

| | hSyn | mDlx |
|---|---|---|
| | Near field neurons | |
| 0.16 $J/cm^2$ | 0.0412 ± 0.0074 | 0.0086 ± 0.0107 |
| 0.30 $J/cm^2$ | 0.0881 ± 0.0084 | 0.0266 ± 0.0126 |
| 0.43 $J/cm^2$ | 0.1770 ± 0.0100 | -0.1183 ± 0.0145 |
| 0.51 $J/cm^2$ | 0.1858 ± 0.0115 | -0.0752 ± 0.0152 |
| 0.61 $J/cm^2$ | 0.1144 ± 0.0173 | -0.0374 ± 0.0133 |
| 0.70 $J/cm^2$ | 0.1720 ± 0.0139 | -0.1606 ± 0.0208 |
| 0.79 $J/cm^2$ | 0.2509 ± 0.0195 | -0.1232 ± 0.0240 |
| | Far field neurons | |
| 0.16 $J/cm^2$ | 0.0214 ± 0.0115 | 0.0362 ± 0.0130 |
| 0.30 $J/cm^2$ | 0.0685 ± 0.0135 | 0.0339 ± 0.0139 |
| 0.43 $J/cm^2$ | 0.1391 ± 0.0168 | -0.0819 ± 0.0167 |
| 0.51 $J/cm^2$ | 0.1529 ± 0.0163 | -0.0473 ± 0.0161 |
| 0.61 $J/cm^2$ | 0.0619 ± 0.0243 | -0.0123 ± 0.0156 |
| 0.70 $J/cm^2$ | 0.1283 ± 0.0213 | -0.1084 ± 0.0272 |
| 0.79 $J/cm^2$ | 0.1706 ± 0.0294 | -0.0918 ± 0.0274 |

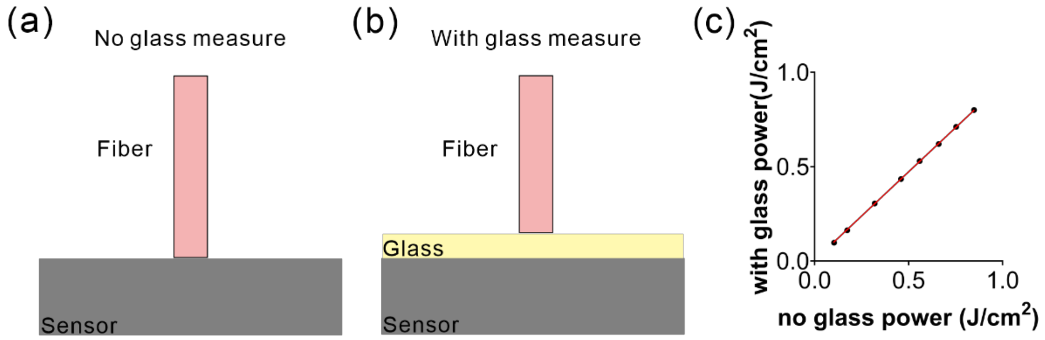


**Fig. S1. Correction of the applied laser energy levels for the glass.**

(a) Setup for laser energy measure with no glass. (b) Setup for laser energy measure with glass. (c) Fitting curve of the correction with glass (red curve, $Y = 0.9363\times X+0.0046$, $R^2 = 0.9998$, $p < 0.0001$). Data represents mean ± SEM.

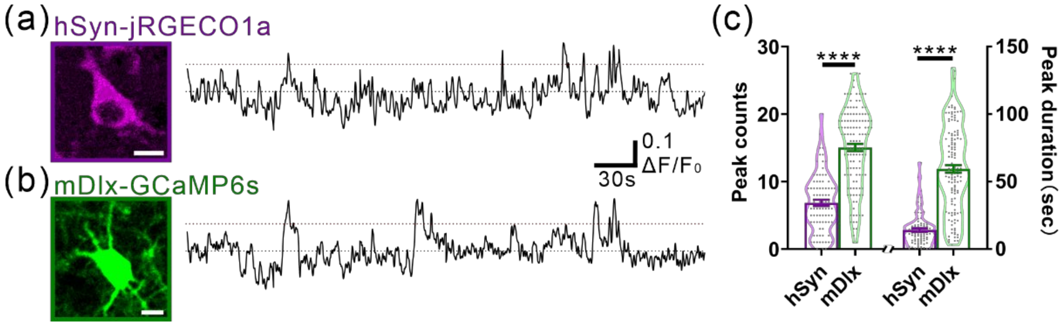


**Fig. S2. Spontaneous calcium activity of hSyn and mDlx neurons.**

(a) Example two-photon in vivo image and consecutive spontaneous activity time course of a hSyn-jRGECO1a labeled neuron. The red dashed line represents the 10% activity threshold value. (b) Example two-photon in vivo image and consecutive spontaneous activity time course of an mDlx-GCaMP6s labeled neuron. (c) Statistical comparisons of peak counts (hSyn: 6.84 ± 0.47; mDlx: 15.05 ± 0.53) and duration (hSyn: 14.07 ± 1.31 s; mDlx: 59.39 ± 2.77 s) of the spontaneous activity between hSyn neurons (n = 83 neurons in 2 mice) and mDlx neurons (n = 129 neurons in 2 mice) (peak counts, $p < 0.0001$; peak duration, $p < 0.0001$). Scale bar, 10 μm; Mann-Whitney test; data represents mean ± SEM.

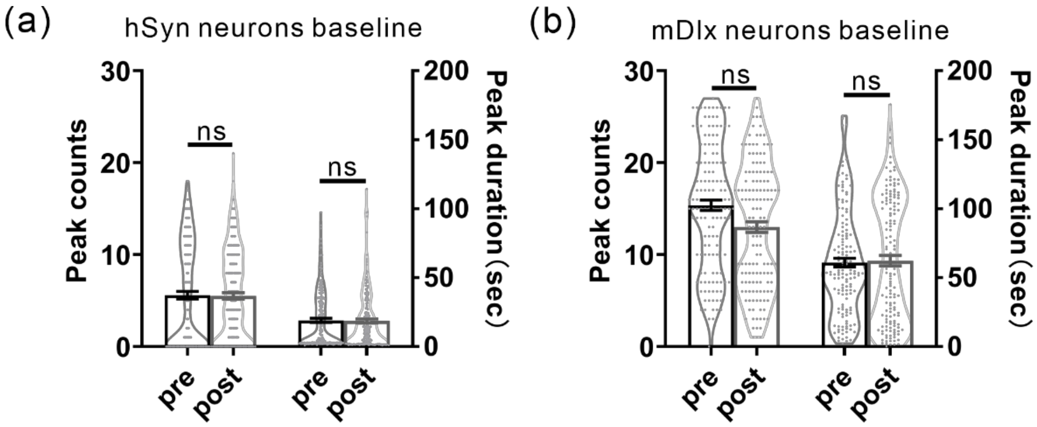


**Fig. S3. No damaging effect on neuronal spontaneous calcium activity post INS stimulation.**

(a) Statistical comparisons of peak counts (pre: 5.59 ± 0.41; post: 5.51 ± 0.35) and duration (pre: 18.97 ± 1.49 s; post: 18.61 ± 1.46 s) of the spontaneous calcium activity between pre-INS and post-INS in same hSyn neuron population (n = 199 neurons in 3 mice) (peak counts, p = 0.5180; peak duration, p = 0.6488). (b) Statistical comparisons of peak counts (pre: 15.37 ± 0.57; post: 13.00 ± 0.57) and duration (pre: 60.87 ± 3.22 s; post: 62.27 ± 3.82 s) of the spontaneous calcium activity between pre-INS and post-INS in same mDlx neuron population (n = 143 neurons in 3 mice) (peak counts, p = 0.0805; peak duration, p = 0.7083). Wilcoxon test; data represents mean ± SEM.

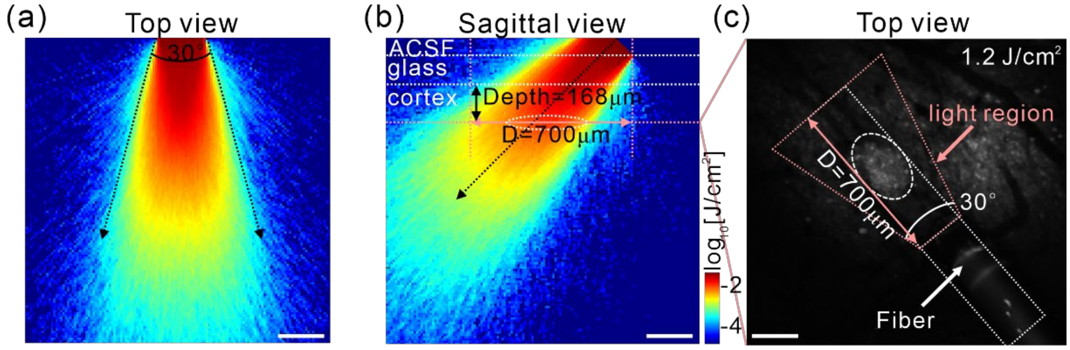


**Fig. S4. Light region definition.**

(a-b) Multi-layer Monte Carlo simulation for infrared light energy. The first layer is ACSF. The second layer is the glass. The third layer is the cortex (imaging depth below cortex: 168 ± 10 μm, from n = 9 mice FOVs). The fiber is at 45° angle. Scale bar, 200 μm. (c) At an averaged structure image, a higher radiant exposure (1.20 J/cm$^2$) is used to guide the center irradiation region and combined with Monte Carlo simulation to define the light region (pink region). Scale bar, 200 μm.

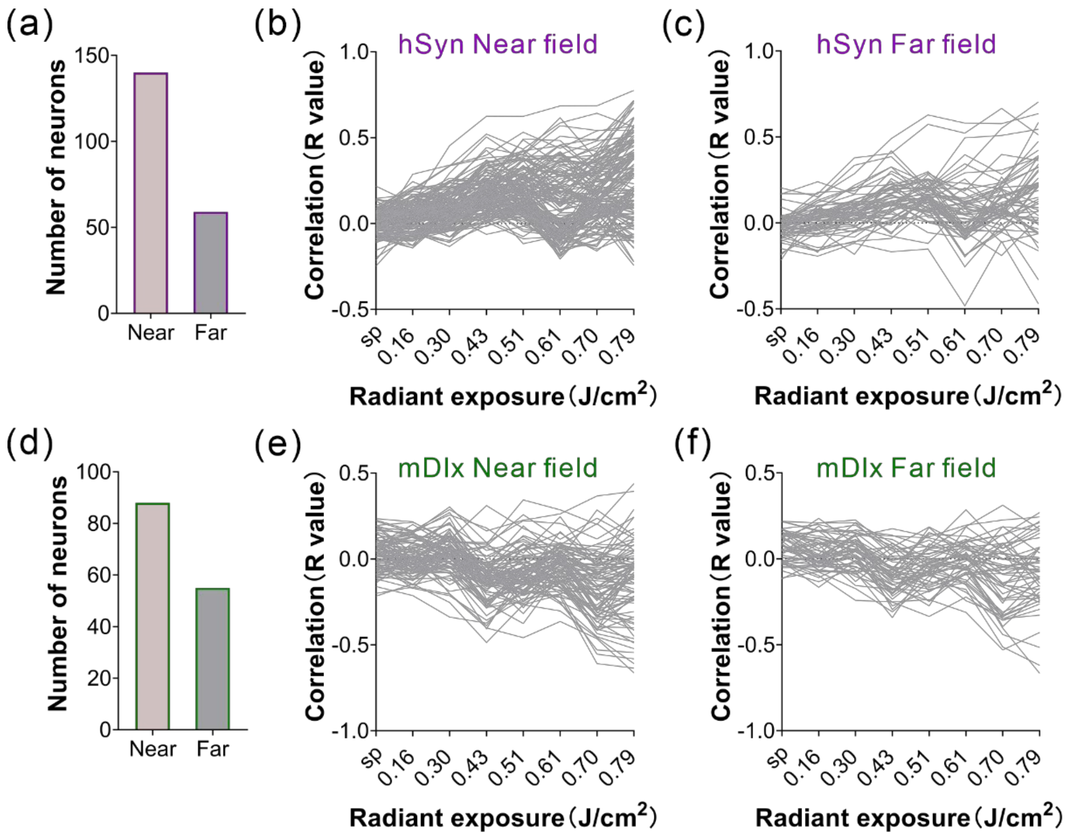


**Fig. S5. Correlation of INS-induced calcium activity of Far field neurons in different subtypes.**

(a-c) Correlation value of hSyn Near and Far field neurons, respectively (n = 140/59 Near/Far field neurons in hSyn mice). (d-f) Correlation value of mDlx Near and Far field neurons, respectively (n = 88/55 Near/Far field neurons in mDlx mice). Data in Table S1.
